# Overcoming High Nanopore Basecaller Error Rates for DNA Storage Via Basecaller-Decoder Integration and Convolutional Codes

**DOI:** 10.1101/2019.12.20.871939

**Authors:** Shubham Chandak, Joachim Neu, Kedar Tatwawadi, Jay Mardia, Billy Lau, Matthew Kubit, Reyna Hulett, Peter Griffin, Mary Wootters, Tsachy Weissman, Hanlee Ji

## Abstract

As magnetization and semiconductor based storage technologies approach their limits, bio-molecules, such as DNA, have been identified as promising media for future storage systems, due to their high storage density (petabytes/gram) and long-term durability (thousands of years). Furthermore, nanopore DNA sequencing enables high-throughput sequencing using devices as small as a USB thumb drive and thus is ideally suited for DNA storage applications. Due to the high insertion/deletion error rates associated with basecalled nanopore reads, current approaches rely heavily on consensus among multiple reads and thus incur very high reading costs. We propose a novel approach which overcomes the high error rates in basecalled sequences by integrating a Viterbi error correction decoder with the basecaller, enabling the decoder to exploit the soft information available in the deep learning based basecaller pipeline. Using convolutional codes for error correction, we experimentally observed 3x lower reading costs than the state-of-the-art techniques at comparable writing costs.

The code, data and Supplementary Material is available at https://github.com/shubhamchandak94/nanopore_dna_storage.

## 1. INTRODUCTION

DNA molecules have been proposed as the storage medium of the future, promising high storage densities (100s of Petabytes per gram [1, 2]) and long-term durability (1000s of years [3]), exceeding the limits of magnetization and solid state storage technologies by several orders of magnitude. DNA storage systems also allow efficient duplication of data and random access using PCR-based techniques [4, 5]. Fig. 1 shows a typical DNA storage system. Binary data is encoded into short DNA sequences (oligonucleotides, or oligos for short) with lengths limited to around 150 bases/nucleotides for current scalable synthesis technologies [6]. Note that each DNA nucleotide belongs to the set {A, C, G, T}. Millions of copies of each oligo are present in the synthesized product, which are then read back using DNA sequencing. The sequencing process involves randomly sampling the oligos, which can lead to duplicated or lost sequences, as well as the loss of ordering among the oligos (permutation). To recover the order of the sequences, typically an index is prepended to each sequence.

**Fig. 1.**
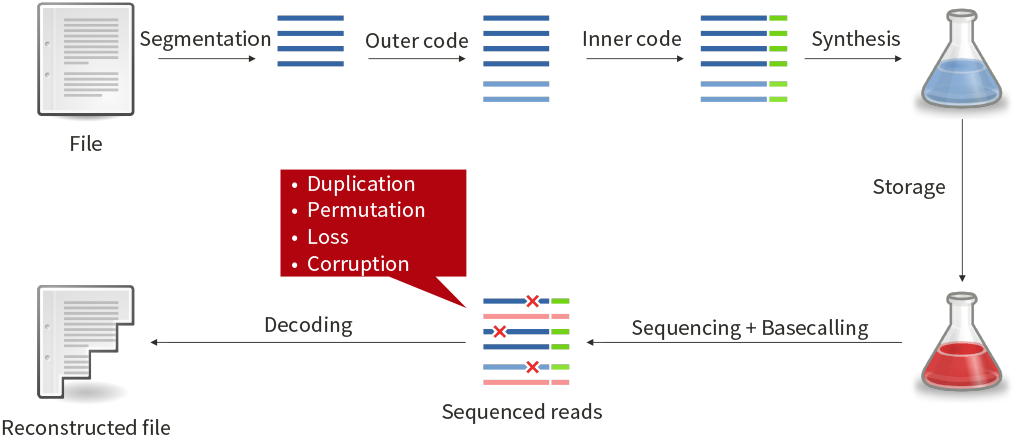
Typical DNA storage system. The input binary file is segmented and encoded using error correction codes before synthesis. The reads obtained after sequencing might have missing sequences or corruptions, which the error correcting decoder seeks to correct.

Moreover, both the synthesis and sequencing processes are noisy and corrupt the sequences with substitutions, insertions and/or deletions of bases. The exact error characteristics are dependent on the specific synthesis and sequencing technology. Successful recovery of the data is facilitated by error correction mechanisms which inject redundancy into the data before synthesis. This typically consists of two components: (i) an outer code (e.g., Reed-Solomon [7] or Raptor [8] code) to recover lost sequences, and (ii) an inner code to detect and/or correct errors within each sequence. Since multiple independent reads can be obtained for each oligo, a consensus operation among the reads can reduce the effective error rate that the inner code needs to handle. This usually involves clustering of reads from the same oligo sequence, followed by a majority vote at each position in the sequence to handle substitutions, and algorithms like trace reconstruction [9] to handle insertions/deletions.

Recent works have examined various aspects of DNA storage, including error correction [1, 5, 10, 11, 12], random access [4, 5, 13], novel synthesis techniques [14, 15] and analysis of the fundamental limits [16, 17, 18]. While initial works used Illumina sequencing which provides highly accurate short reads, there is growing interest in the use of nanopore sequencing [19] because it is a portable, real-time and low-cost platform that also supports long reads. The nanopore sequencer first outputs a current signal induced by the DNA sequence, which is then translated back to a sequence by the basecaller. Nanopore sequencing poses significant challenges due to higher error rates in the basecalled reads (10% as compared to 1% for Illumina), including a substantial number of insertions and deletions, which are notoriously difficult to correct [20]. Due to the high error rate and the lack of suitable inner codes for insertion/deletion errors, most previous works [5, 14, 21, 22] rely heavily on consensus algorithms working with a large number of reads, leading to very high reading costs. This is due to the similarity of consensus with repetition coding, which is known to be suboptimal. One strategy to avoid the high basecalling error rate and consensus could be to work with the raw current signal which has much higher information content than the basecalled read. Unfortunately, the raw signal is hard to model and difficult to use directly in error correction decoding.

In this work, we propose a novel basecaller-decoder integration approach for nanopore sequencing-based DNA storage, in which the decoding is performed on the intermediate output of the basecaller, rather than the final basecalled reads which exhibit high error rates. This intermediate output is more informative about the raw signal than the basecalled reads, while also being easier to model and integrate with error correction schemes than the raw signal itself. To match the sequential Markov nature of the nanopore sequencing and basecalling process, we use a convolutional coding scheme [23] as the inner code and achieve 3x lower reading cost than the state-of-the-art works at similar writing cost.

## 2. NANOPORE SEQUENCING AND BASECALLING

In this section we briefly describe nanopore sequencing and basecalling to motivate our proposed approach. Nanopore sequencing, in particular the MinION sequencer by Oxford Nanopore Technologies (ONT) [19], offers a portable and real-time solution capable of sequencing long reads. The sequencing process involves a single strand of DNA passing through a pore leading to variations in a measured ionic current. The current at any given time primarily depends on a nucleotide subsequence of length 6 (i.e., a 6-mer) inside the pore at that instant. This raw current signal is sampled (typically at a frequency of 4 kHz), and is used by the basecaller to infer the most likely base sequence that could have induced the raw signal.

**Fig. 2.**
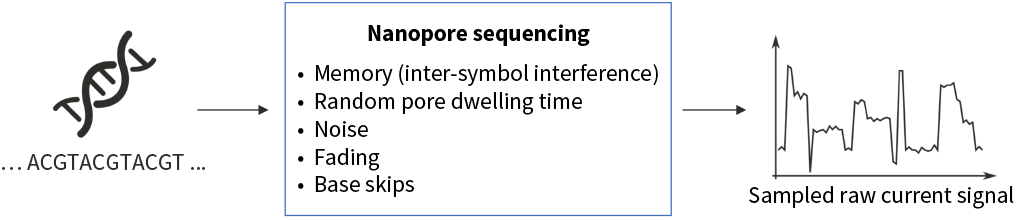
Signal distortions caused by nanopore sequencing, viewed from the channel coding perspective.

Fig. 2 shows the various causes of distortions in the nanopore sequencing process from the channel coding perspective [24, 25]. Firstly, as the raw signal depends on multiple bases (usually 6 bases) at a time, this causes inter-symbol interference. The “pore dwelling time”, i.e., the number of signal samples per 6-mer, randomly varies between 5-15 samples, leading to synchronization issues. Each sample for a given 6-mer undergoes noise and fading (variation in mean current value for a given 6-mer). In some cases, the pore dwelling time can be 0, leading to base skips.

Due to the challenges associated with modeling the nanopore sequencing process, there has been a shift from analytical modeling to deep learning based modeling in the state-of-the-art basecallers [26, 27]. For instance, the Flappie basecaller [28] by ONT uses a recurrent neural network to translate the raw current signal into probabilities of transition between pairs of bases at each time step (see Supplementary Material for the network architecture). The most likely base sequence is then determined from these probabilities using the Viterbi algorithm [29] (Fig. 3(a)). A similar approach is used in other basecallers such as Chiron [30] and Scrappie [31]. Despite the advances due to deep learning based basecalling, the error rate of basecalled nanopore reads is still around 10%, with a significant fraction of insertion and deletion errors [27].

**Fig. 3.**
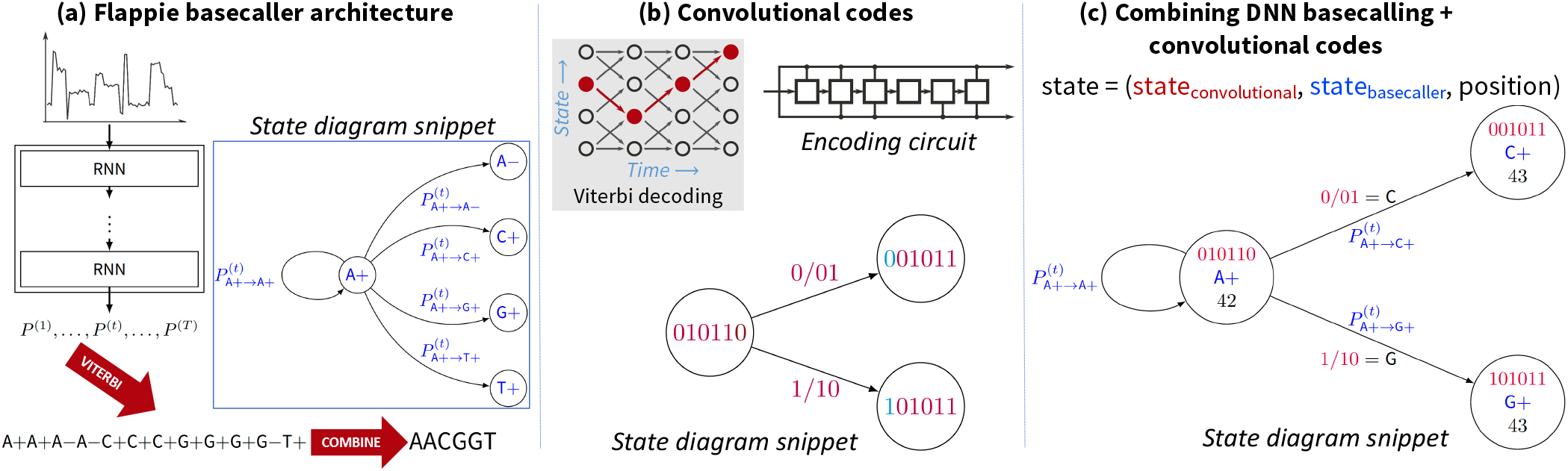
Integrating basecaller and convolutional code decoder. (a) The architecture of the Flappie basecaller, which consists of a recurrent neural network (RNN) that predicts the probabilities of base transitions at each time step 1, …, *T* in the raw signal which are then fed into a Viterbi decoder to obtain the most likely base sequence. To distinguish between “stay transitions” (i.e., no movement of DNA) and transitions due to repeated bases (e.g., AA), two states (+, −) are used per base and a collapsing operation is performed to obtain the basecalled sequence. (b) The encoding circuit for a rate 1/2 convolutional code with a shift register storing the current state, a snippet of the state transition diagram, and the Viterbi decoding process illustrated as a trellis with the optimal path highlighted. (c) Basecaller-decoder integration by combining the convolutional code state with the basecaller state, producing the most likely path that forms a valid codeword. A snippet of the combined state transition diagram is shown. The state also includes the position in the codeword to ensure correct length of the optimal path.

Unlike typical biological applications, DNA storage provides us with the flexibility to design the DNA sequences and knowledge of this additional structure can be leveraged in basecalling. Instead of the approach shown in Fig. 1 where the inner code decoder operates on the basecalled sequence, we propose integrating the decoding with the basecalling, thus taking into account the inner code structure. This is achieved by applying the decoding on the intermediate transition probabilities from the recurrent neural network in the Flappie basecaller. By working with the transition probabilities, our approach sidesteps the difficulties of working directly with the raw current signal, while still utilizing much of the rich soft information in the raw signal as distilled by the neural network.

## 3. METHODS

We now describe the encoding and decoding architectures in our proposed approach. We follow the framework shown in Fig. 1 for the most part, using a Reed Solomon code [7] as the outer code and a convolutional code [23] along with CRC error detection [32] as the inner code. The major difference is that the basecaller and the inner code decoder are now integrated, allowing us to utilize the rich soft-information in the raw signal. We next describe the individual components of our system followed by a brief discussion of the complete framework. Details regarding the parameters and the implementation are available in the Supplementary Material.

### Convolutional code as the inner code

In convolutional codes, the encoder encodes a stream of message bits into a sequence of encoded bits, which are computed as a linear combination (over 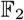) of a past window of *m* input bits (*m* denotes the memory of the code). At any given time step, the value of the past m input bits denotes the “state” of the convolutional code, e.g., Fig. 3(b) shows a snippet of the state transition diagram (showing the states along with the input/output on the transitions) for a convolutional code with *m* = 6 and rate 1/2, producing 2 output bits per input bit. As *m* increases, the code becomes more powerful, while the decoding becomes slower due to an exponential increase in the number of states with *m*.

We use rate 1/2 convolutional codes with *m* = 8, 11, 14 that were among the best short block codes studied in [33]. To achieve rates higher than 1/2, we use the technique of puncturing, which removes a pre-defined pattern of bits from the output [34]. As seen below, the sequential structure of convolutional codes make them a natural fit for integration with the basecalling process for the nanopore raw current signal which can be modeled as a sequential hidden Markov process [26]. Convolutional codes also provide an elegant method to handle reverse complemented reads based on the fact that the reverse of a convolutional code is also a convolutional code with similar error correction capabilities.

### Basecaller-decoder integration

The decoding of the convolutional code is performed using the transition probabilities generated by the basecaller, rather than the final basecalled sequence. As shown in Fig. 3, a combined state is formed from the state of the basecaller, the state of the convolutional code and the position in the codeword. The code structure dictates the possible transitions while the probabilities from the basecaller are used for scoring possible paths. The most likely path is obtained using the Viterbi algorithm [29] (dynamic programming) and the corresponding input message bits are obtained based on its state transitions.

### List decoding and CRC

In spite of the improved inner code decoding error rate due to the proposed approach, there remains a sizable fraction of incorrectly decoded reads that are difficult to handle with the outer code alone. Therefore, we add a 8-bit cyclic redundancy check (CRC) to the the payload and index (Fig. 4(a)), which allows detection and removal of erroneously decoded sequences. Note that for an 8-bit CRC, there is a 1/256 probability of an erroneous sequence satisfying the check, and hence the outer code needs to be designed to handle a small number of errors.

**Fig. 4.**
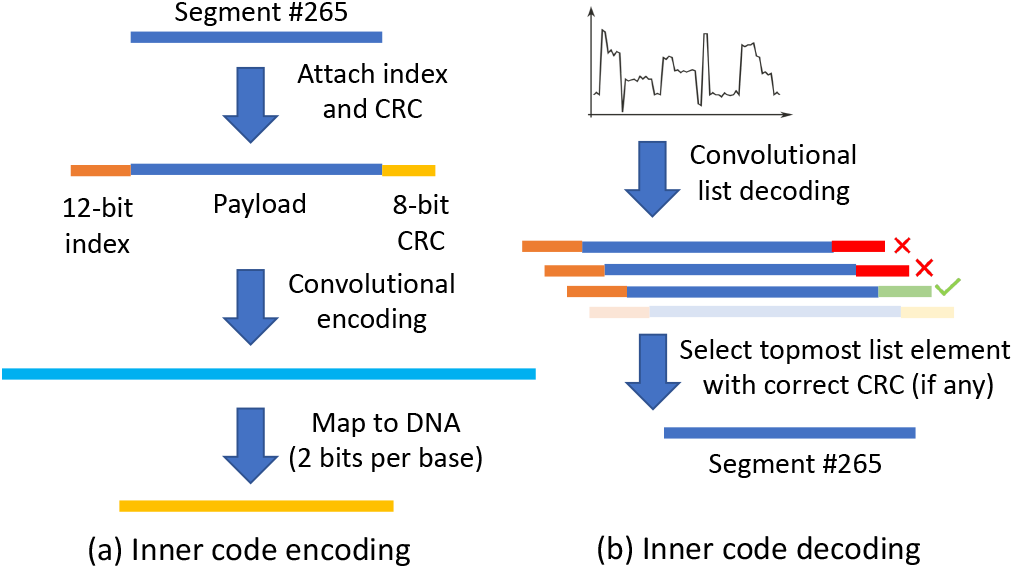
Inner code encoding and decoding. (a) Index and CRC are added to each segment produced by the outer code. This is followed by convolutional encoding and mapping to the DNA alphabet. (b) During decoding, the raw signal is decoded into a list of most likely codewords by the Viterbi decoder. The topmost element in the list that satisfies the CRC (if any) is used for outer code decoding.

The use of the CRC error detection also enables list decoding for the convolutional code, wherein a list of the *L* most likely input message sequences are generated by the Viterbi decoder. Out of these, the topmost element satisfying the CRC is chosen (if any). The list decoding is based on [35] with modifications made to account for the possibility of multiple state sequences leading to the same input message due to the stay transitions (see Fig. 3(c)).

### Reed-Solomon outer code

We use a Reed-Solomon (RS) code with field size 2^16^ as the outer code to recover lost sequences and to correct any errors left undetected by the CRC. The amount of additional RS redundancy can be chosen to tradeoff the writing and reading costs [12], and is set to 30% by default. Our RS implementation is similar to that in [5] and is based on the Schifra library [36].

Fig. 4 shows the overall inner code encoding and decoding procedure. During encoding, an index is prepended to each segment produced by the outer RS code and then a CRC of the index and payload is appended to it. This is encoded with the convolutional code and then mapped to DNA using a 2 bits/base encoding (00→A, 01→C, 10→G, 11→T). The decoding process, as discussed above, involves list decoding of the convolutional code, selection of the message from the list based on the CRC and outer RS code decoding.

## 4. EXPERIMENTAL SETUP

We evaluated the proposed scheme by performing experiments with various settings of the parameters *m* (convolutional code memory), *r* (convolutional code rate) and outer RS code redundancy. For each experiment, the same file was encoded into DNA sequences of length ≈ 115. The sequences were synthesized by Customarray (http://www.customarrayinc.com/) in a single pool of 12K oligos after adding primer sequences of length 25 on both sides, leading to a total oligo length ≈ 165. The primers are required for PCR amplification before sequencing. The encoded file of size 11 KB consisted of several texts including the UN declaration of human rights, the Gettysburg Address, the “I Have a Dream” speech, a set of poems and the lyrics of Rick Astley’s “Never Gonna Give You Up”. The files were compressed and encrypted before encoding so that the input to the encoder appears random and does not produce excessive homopolymers (e.g., GGGG) which cause synthesis and sequencing errors [12]. The synthesized pool of oligos was amplified with PCR and sequenced using an ONT MinION sequencer (R9.4.1 pore). For convenience, the raw signals corresponding to each parameter experiment were separated computationally. In practice, this can be achieved using PCR-based random access [5] using the experiment-specific primers.

For each experiment, the minimum number of reads required for successful decoding was obtained. To ensure robustness, random subsampling of reads was performed and the reported results indicate success in 10 out of 10 such trials. We used writing cost (measured in bases synthesized per information bit) and reading cost (measured in bases sequenced per information bit) as the figures of merit (excluding the primers for both quantities). A recent work [12] showed a tradeoff between these quantities in the context of Illumina sequencing based DNA storage and recommended the use of reading cost as the metric rather than coverage (measured in bases sequenced per bases synthesized). We compare our results to previous works [5, 22]. While there have been other works on DNA storage using nanopore sequencing [14, 21], their results are not included in the comparison due to the use of significantly different synthesis strategies, different pore versions (R7 vs. R9.4.1) and/or limited information regarding the minimum reading cost achieved.

We note that a direct comparison across works is difficult due to the use of different synthesis providers, different amounts of encoded data and different oligo lengths. Furthermore, [5] and [22] use MinION’s 1D^2^ sequencing mode which has lower error rates (7%) and lower throughput as compared to 1D sequencing mode used in this work. Despite this, the comparison is useful in providing an estimate of the benefits of the proposed strategy.

More details about the experiments, including the removal of the primers for convolutional code decoding and the handling of reverse complemented reads, are available in the Supplementary Material at https://github.com/shubhamchandak94/nanopore_dna_storage.

Fig. 5 shows the tradeoff between writing and reading costs achieved by the proposed scheme and the previous works. For the proposed scheme, it shows results for convolutional codes with memory *m* = 8, 11, 14, each with three rates *r* = 1/2, 3/4, 5/6. Comparing the *m* = 11, *r* = 5/6 code to the previous state-of-the-art [5, 22], we observe a 3x improvement in the reading cost at the same writing cost. This suggests that the basecaller-decoder integration approach can lead to significant improvements over the traditional approach of applying the decoder on the basecalled reads with heavy reliance on consensus. Our work brings the read/write cost tradeoff achieved for nanopore based DNA storage closer to the state-of-the-art for Illumina based DNA storage [12]. In particular, we require only 2x higher reading cost, despite the former having an order of magnitude higher basecalling error rate than the latter.

**Fig. 5.**
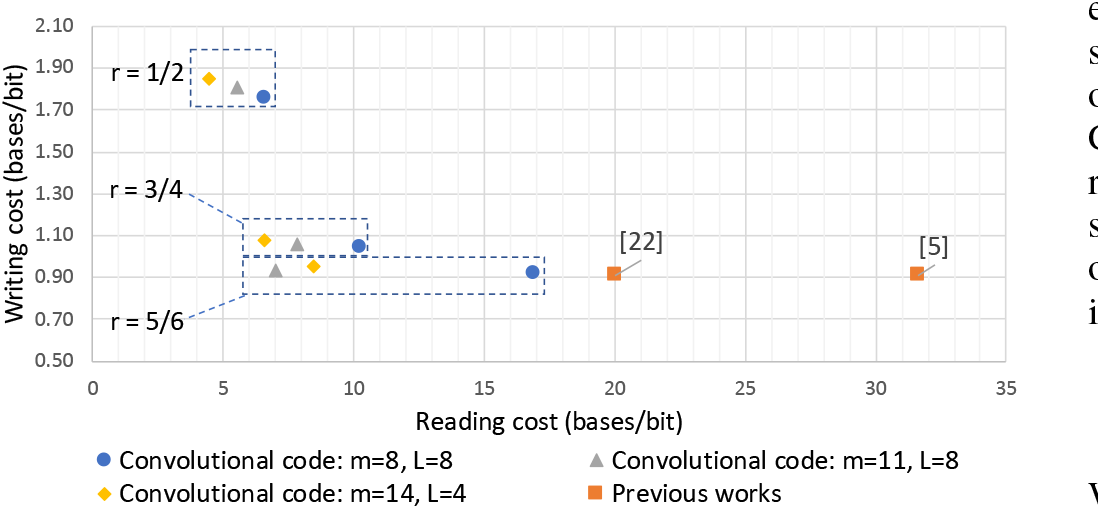
Comparison of writing and reading cost achieved by the proposed scheme and previous works. Multiple parameters (convolutional code memory m, rate r, list size L) were considered, achieving 3x lower reading cost than the previous state-of-the-art at the same writing cost. Note that the writing and reading cost are bounded below by 0.5 since each base can represent at most 2 bits.

Comparing the results for a fixed memory *m* of the convolutional code, as we increase the rate *r*, we observe lower writing cost and higher reading cost. This is because the error correction capability of the code decreases with increasing *r*. As *m* increases, we observe lower reading cost at the same writing cost, at the cost of slower decoding, with diminishing returns beyond *m* = 11.

The results for the proposed scheme shown in Fig. 5 were obtained with list size *L* = 8 for *m* = 8, 11. List size *L* = 4 was used for *m* = 14 due to computational constraints. Recall that the convolutional code decoder produces a list of *L* candidate sequences, out of which the topmost candidate satisfying the CRC is chosen. As the list size increases, we observe a reduction in the reading cost because of an increase in the probability of the true sequence being part of the list. However, at very high list sizes (≈ 32), we observe an inversion in this trend. This is due to an increase in the probability of an incorrect sequence satisfying the CRC appearing in the list. This probability can become significant due to the short CRC length of 8 bits (chosen to minimize the writing cost), leading to higher reading costs since more of the outer code redundancy is used up for error correction rather than for the recovery of missing oligos. The fraction of reads for which the correct sequence was present in the list varied from 20% to 80% depending on the values of *m, r* and *L*. In particular, for *m* = 11, *r* = 5/6, *L* = 8, this fraction was 25%, suggesting that on an average only 4 copies of an oligo were required to decode it successfully. More detailed results on various parameters including list size and outer code redundancy are available in the Supplementary Material.

Computationally, the convolutional code decoding is the most expensive step in the proposed decoding algorithm, with time and space complexity 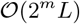 where *m* is the convolutional code memory and *L* is the list size. For the current implementation written in C++, decoding a single read for (*m* = 11, *r* = 5/6, *L* = 8) took roughly 270 s with RAM consumption of 1.3 GB (Ubuntu 18.04.1 server with 2.2 GHz Intel Xeon processor, single-threaded). As part of future work, we plan to explore algorithmic and implementation improvements to speed up the decoding.

## 6. CONCLUSIONS AND FUTURE WORK

We proposed a novel basecaller-decoder integration approach for nanopore sequencing based DNA storage which achieves 3x lower reading cost compared to earlier works at similar writing cost. Instead of working with the basecalled reads which exhibit high error rates, the proposed decoder utilizes the soft information available in the basecaller pipeline. We use convolutional coding as the inner code, with its sequential Viterbi decoding integrated with the nanopore basecaller architecture. Future work includes optimization of convolutional code parameters, finetuning of neural network parameters in the basecaller, improvements in computational efficiency using approximate decoding and combining the strengths of this framework with those of consensus-based techniques.

The basecaller-decoder integration approach is useful beyond convolutional codes and can also be applied to novel synthesis techniques such as enzymatic synthesis [14] and k-mer-by-k-mer synthesis [37]. It can also be useful for utilizing the information in the raw current signal for bioinformatics applications beyond DNA storage.

Finally, we believe that this hybrid approach of combining ML-based modeling with analytical error correction coding schemes can be extended to other communication and storage problems. This is a natural approach towards combining recent advances in machine learning with the significant research done in information theory and communication.

