## Supplementary Material for "Overcoming High Nanopore Basecaller Error Rates for DNA Storage Via Basecaller-Decoder Integration and Convolutional Codes"

**3 Reed Solomon outer code** 5

**4 Simulations** 6

**5 Experimental parameters and results** 6

**6 Experimental procedure** 10

**7 Basecalling analysis** 10

#### 1 Availability of code and data

The code and instructions for installation and encoding/decoding are available at `https://github.com/shubhamchandak94/nanopore_DNA_storage/`. The specific scripts used for running various experiments are mentioned in the README and the text below as relevant. The data (encoded files, oligo files sent for synthesis, raw sequencing data, error statistics, decoded lists) is available at `https://github.com/shubhamchandak94/nanopore_DNA_storage_data/`.

### 2 Inner coding

#### 2.1 Convolutional code parameters

We use three rate 1/2 convolutional codes with memory $m = 8, 11, 14$ for our experiments. The code parameters were derived from [1] and are given below. The current implementation also supports another code with $m = 6$ which is obtained from [2]. Note that a rate 1/2 convolutional code can be specified using the two output functions, which represent the linear combination of the state and current input used to obtain the output. Following the convention in [1], the output linear combinations (i.e., generator polynomials) are written in octal.

- $m = 6$: $G = [171, 133]$
- $m = 8$: $G = [515, 677]$
- $m = 11$: $G = [5537, 6131]$
- $m = 14$: $G = [75063, 56711]$

To better relate the parameters to the encoding circuit for $m = 6$, note that $G$ can be written in binary as $[1111001, 1011011]$ where the 1's show the position of the connections in the encoding circuit in Figure 1. Note that the two output streams are interleaved, i.e., if the first output stream (shown as the top output) is $(C_1(1), C_1(2), C_1(3), \dots)$ and the second output stream (shown as the bottom output) is $(C_2(1), C_2(2), C_2(3), \dots)$, then the overall output is $(C_1(1), C_2(1), C_1(2), C_2(2), C_1(3), C_2(3) \dots)$

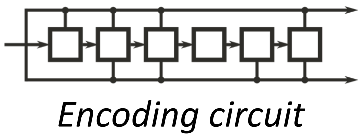

Figure 1: Encoding circuit for $m = 6$ code with $G = [171, 133]$.

#### 2.2 Start/end convolutional code state and impact of runs

The convolutional code encoding typically starts and ends with a fixed and known state (usually the all zeros state) [3]. This is achieved by padding the input with $m$ known bits at the end, thus guaranteeing a specific end state. For this work, we found that starting/ending with the all zero state was suboptimal because of synchronization issues. For example if the input bit sequence is 000111...1001011, the sequence of states starting from the all zeros state (for $m = 6$) would be $000000, 000000, 000000, 000000, 100000, 110000, \dots$. Due to the repeated sequence of states at the beginning, the decoder tends to incorrectly predict the number of 0s at the start. This is because the number of raw signal samples corresponding to each base is random, a fact that leads to difficulties in inferring the length of repeated state sequences. The result is that the decoder tends to produce a shifted message. To resolve this issue, we adopt start and end states that do not lead to repeated states. The start states used are listed below (the end states were chosen as the reverse of the start states for symmetry in case of reverse complemented reads).

- $m = 6$: 100101
- $m = 8$: 10010110
- $m = 11$: 10010110001
- $m = 14$: 10010110001101

#### 2.3 Convolutional code puncturing for higher rates

To increase the convolutional code rate beyond 1/2, we use the technique of puncturing [2]. The idea is to transmit only a prespecified subsequence of the bits, usually dictated by a repeating pattern. For example, suppose that $C_1(i)$ and $C_2(i)$ represent the two output streams of the code. Then the output without puncturing is given by $(C_1(1), C_2(1), C_1(2), C_2(2), C_1(3), C_2(3) \dots)$. Now, if we use a puncturing pattern
$C_1$ : 0 1
$C_2$ : 1 0
where 1 represents a transmitted symbol and 0 represents a non-transmitted symbol, then the output would look like $(C_2(1), C_1(2), C_2(3), C_1(4) \dots)$ and the new code rate is 1, i.e., one output bit per input bit. The current implementation uses the following patterns (generally based on [2] with some modifications due to reasons described below).

- Rate = 1/2
  $C_1$ : 1
  $C_2$ : 1
- Rate = 2/3
  $C_1$ : 1 1 0 1
  $C_2$ : 1 0 1 1
- Rate = 3/4
  $C_1$ : 1 0 1
  $C_2$ : 1 1 0
- Rate = 4/5
  $C_1$ : 1 0 0 1 1 0 0 1
  $C_2$ : 1 1 1 1 0 1 1 0
- Rate = 5/6
  $C_1$ : 1 0 1 1 0
  $C_2$ : 1 1 0 0 1
- Rate = 7/8
  $C_1$ : 1 0 0 0 1 0 1
  $C_2$ : 1 1 1 1 0 1 0

The binary output is converted to a DNA sequence with a 2 bits per base mapping (00→A, 01→C, 10→G, 11→T). During the decoding the state transitions of the basecaller (Flappie) correspond to either a stay (no change in current base) or a shift by one base. When the convolutional code rate is 1/2, each basecaller transition by one base corresponds to one state transition in the convolutional code (since each input bit produces one base). With puncturing, the situation is a bit more involved. To simplify things as much as possible, we only use puncturing patterns composed of the following building blocks:

$$\begin{matrix} 1 & & 0 & 1 & & 1 & 0 & & 0 & 0 & & 1 & 1 \\ 1 & , & 1 & 0 & , & 0 & 1 & , & 1 & 1 & , & 0 & 0 \end{matrix}$$

In each of these cases, either one or two input bits produce two output bits (i.e., one base). When one input bit produces one base, an example state transition in the decoder would be from $(010011, A+, 42)$ to $(001001, C+, 43)$ where the convolutional code state shifts by one. In this case, there are three transitions out of the state (one stay, and two corresponding to input bit 0 or 1). When two input bits produce one base, an example state transition in the decoder would be from $(010011, A+, 42)$ to $(000100, C+, 43)$ where the convolutional code state shifts by two. In this case, there are five transitions out of the state (one stay, and four corresponding to pairs of input bits). Finally, note that in some cases the input needs to be padded by one bit to make sure that the puncturing pattern is composed of the given building blocks and the total number of output bits is even. For example, in the case of Rate 7/8, if the message + termination length is 2 mod 7, then one padding bit is added so that we have even number of output bits.

#### 2.4 Handling reverse complemented reads

The sequencing process involves double stranded DNA where the sequence or its reverse complement can be sequenced. The reverse complement sequence consists of the original sequence in reverse, where each base is replaced by its complement (A↔T, C↔G). The decoding of reverse complemented reads is based on the fact that reversing a codeword for a convolutional code produces a codeword for a different convolutional code which has the same minimum distance properties but different generator polynomials. In fact, the generator polynomials and the puncturing patterns are effectively reversed for the new convolutional code. For example, if the generator polynomial $G$ written in binary is $[1111001, 1011011]$, the corresponding polynomial for reverse complement reads is $[1001111, 1101101]$. The output mapping from bits to bases in the reversed code is also modified appropriately. Thus, the handling of reverse complemented reads is quite straightforward as long as we can detect whether the read is reverse complemented. The detection is done based on the PCR primers as discussed in the next section.

#### 2.5 Removal of primers for convolutional code decoding

The raw current signal obtained from the nanopore corresponds to a DNA sequence consisting of the PCR primers and sequencing adapters in addition to the encoded sequence. The primers are known sequences of length 25, and are required for PCR amplification before sequencing. As the Viterbi decoder only expects the transition probability array corresponding to the encoded sequence, it needs to be trimmed appropriately. To identify the part corresponding to the encoded sequence, we first perform basecalling using Flappie. Then we search for the start and end primer in the basecalled sequence based on minimum edit distance. Based on the position of the primers, we truncate the transition probability array that the decoder receives. To detect reverse complemented reads, we perform the minimum edit search for the primers in both orientations (forward and reverse complemented) and pick the one that achieves lower edit distance.

#### 2.6 Flappie basecaller architecture

The Flappie basecaller [4] consists of a recurrent neural network followed by Viterbi decoding and a collapse operation. The pretrained recurrent neural network model provided by ONT has the following structure:

```
Convolution-layer(stride=2, filter-size=19, num-filters=256)
Tanh Nonlinearity
Reverse-GRU Layer
GRU Layer
Reverse-GRU Layer
GRU Layer
Reverse-GRU Layer
Fully-Connected output layer
```

The network consists of a 1D-convolution layer, and 5-layer GRU network (with alternating Forward and Reverse layers). The neural network takes as input the raw signal and outputs a transition probability matrix at each time step in raw signal (more precisely, the raw signal is downsampled by 2 with the initial convolutional layer with stride 2). The transition probability matrix contains the posterior probability of each allowed state transition at the given time step given the raw signal data. From this data, the Viterbi decoding produces the most likely sequence of states (A+, A−, C+, C−, G+, G−, T+, T−) given the observed raw signals, based on a conditional random field (CRF) framework [5]. Runs of same states are then collapsed to produce the final basecalled sequence, for example, a state sequence A+A+A+A−A−A−C+C+C+C+T+T+G+G+G+G− is collapsed to AACTGG. Note that using two states (+ and −) per base allows us to distinguish between repeated bases in the sequence (e.g., A+A−) and multiple time steps (i.e., raw signal samples) corresponding to the same base (e.g., A+A+).

#### 2.7 List Viterbi decoding and implementation details

For the Viterbi decoding process, the state includes the convolutional code state, the Flappie basecaller state and the position in the codeword. The most likely state sequence that ends in a valid state is obtained using

Viterbi decoding. The valid end state must have the correct convolutional code end state (see Section 2.2) and the position equal to the codeword length. When the position is not included in the state, we found that the optimal state sequences produced significantly shorter message lengths, thus we decided to include the position in the state to enforce the correct message length in the decoded output. The scores for the transitions are obtained from the transition probability matrix from the basecaller.

Note that the total number of states is given by the product of (i) the number of convolutional code states ($2^m$ where $m$ is the memory of the code), (ii), number of Flappie basecaller states (8: A+, A−, C+, C−, G+, G−, T+, T−), (iii) message length (approx. 100-200 depending on code rate and oligo size). For $m = 11$, the total number of states is more than 1 million, which can lead to computational issues during decoding as described below. Despite the large number of states, the transitions are very sparse since only transitions from the same or previous codeword position are allowed into a given state, and the convolutional code state and basecaller states have their own transition constraints. In particular, for the basecaller state, the allowed outgoing transitions from A+ or A− are to the states A+, A−, C+, G+, T+ (similarly for other bases).

To further improve the error rates over usual Viterbi decoding, we perform list decoding instead of ML decoding, obtaining the top $L$ most likely messages rather than just the most likely message. Out of these, the topmost element satisfying the CRC is chosen (if any). We use a 8-bit CRC, where the short CRC length was chosen to reduce the writing and reading cost. The crc8 library in Python3 (`https://pypi.org/project/crc8/`) was used for the implementation. In cases where we get multiple reads with the same index that satisfy the CRC, we use the most commonly occurring sequence among those for the outer code decoding. Note that the final block error rates are relatively high (∼20%-80% depending on parameters, see results). Thus, the CRC plays a critical role for error detection and successful outer code decoding.

We use the parallel List Viterbi Algorithm from [6] with certain modifications. The list algorithm attempts to find the $L$ most likely paths (i.e., sequences of states) instead of just the most likely path. This is done by keeping track of the top $L$ paths entering each state at each time step. We initially faced two major issues with this: (i) since multiple state sequences can correspond to the same message (due to stay transitions), finding the top $L$ best paths through the states tends to produce a list containing just 1 or 2 distinct messages (rather than $L$), (ii) the memory consumption grows linearly with $L$ and rapidly becomes intractable, chiefly due to the traceback array which needs to store $L$ entries (previous state) for each state at each time step.

To resolve these issues, we utilize the fact that we do not need to know the most likely state sequence, instead we are interested in the messages themselves. Thus, instead of using a traceback array that needs to store the previous state corresponding to the best path at each state at *each* time step, we just store a list of $L$ most likely messages along with their scores at each state at the *current* time step. Storing the messages at the current state also allows us to keep only the distinct messages at each state. The implementation uses heaps to perform a merge of the incoming lists at each state to find out the list of $L$ most likely distinct messages at this state.

Finally, to further optimize the decoding, we do not allow states for which the position in the codeword deviates significantly from the expected position based on the location in the raw signal. Allowing a large enough deviation (±20 time steps) still provides significant speedup while having essentially no impact on the optimality. The implementation is parallelized and can operate on the different states at a given timestep in parallel. If sufficient memory is available, further speedup can be gained by running the decoder on different reads in parallel.

### 3 Reed Solomon outer code

We use Reed Solomon (RS) code as the outer code to recover lost sequences and to correct any erroneous sequences which were not detected by the CRC. The RS code implementation is similar to that in [7] and is built on the Schifra library [8].

Figure 2 illustrates the Reed Solomon encoding procedure. The input data is segmented into segments of length $16n_{RS}$ where $n_{RS}$ is an integer. The length of the segment is decided based on the desired length of the synthesized oligos after the inner coding. The RS coding with symbol size of 16 bits is applied independently

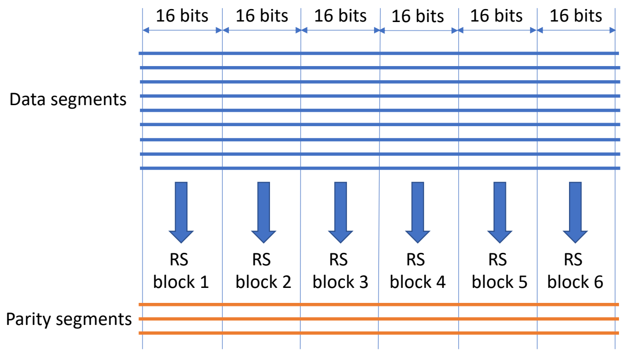

Figure 2: Reed Solomon encoding schematic.

to the $n_{RS}$ chunks of the segments as shown in the figure, generating new parity segments. If the number of data segments is $n_{data}$ and the RS redundancy is $f$ (30% by default), then the number of parity segments $n_{parity} = fn_{data}$. Since we are working with symbol size of 16 bits, $n_{data} + n_{parity} < 65536$, which means that $n_{data} < n_{data}^{MAX} := 65536/(1+f)$. For larger data sizes, we need to apply the procedure independently to groups of $n_{data}^{MAX}$ segments.

Smaller symbol sizes lead to very few segments in each block, leading to high probability that some of blocks fail in the decoding (due to channel dispersion effects). Larger symbol sizes lead to higher computational complexity and also lack good existing implementations. The choice of 16 as the symbol size was made to tradeoff these factors. To understand why RS coding is applied to vertical blocks as shown in Figure 2, note that the erasures always occur at the segment-level. The vertical strategy allows erasure coding over a larger number of segments (up to 65,535 including parity segments) and similar strategies are often applied for bursty errors (interleaving).

During decoding, the set of segments decoded by the inner code are used as input to the RS code decoder. For each block, the decoding succeeds as long as $n_{erasure} + 2n_{error} < n_{parity}$ where $n_{erasure}$ is the number of erased (lost) segments and $n_{error}$ is the number of erroneous segments (due to CRC detection failure). An increase in $n_{error}$ leads to the RS code being able to handle fewer erasures which significantly increases the reading cost, especially in the presence of coverage bias [9]. Thus, a sufficiently long CRC is required for successful RS code decoding.

### 4 Simulations

We provide an end-to-end simulator for the inner convolutional code (code: `simulator.py`). The simulator supports features such as synthesis errors (substitutions, insertions, deletions), reverse complementation, and nanopore raw signal generation. For the nanopore raw signal simulation, we tried two simulators, scrappie [10] and DeepSimulator [11]. Overall, we found that scrappie provided results closer to what we observed in practice, even though it's dwell time is quite idealistic and has low variance. Therefore, we provide an option to use the dwell times from scrappie or DeepSimulator while using the signal levels (mean and variance) from scrappie. Several parameters for the convolutional code including the CRC size, convolutional code memory, puncturing pattern, code rates and list size were chosen based on this simulator.

### 5 Experimental parameters and results

#### 5.1 Encoded files

The input file used for our experiments consisted of the following text files:

- gettysburg.txt: The Gettysburg Address by Abraham Lincoln
- mlkjr.txt: "I have a Dream" by Martin Luther King, Jr

- poems.txt: A collection of poems:
  - “The Road Not Taken” by Robert Frost
  - “Stopping by Woods on a Snowy Evening” by Robert Frost
  - “If” by Rudyard Kipling
  - “Fire and Ice” by Robert Frost
  - “The Tyger” by William Blake
  - “A Psalm of Life” by Henry Wadsworth Longfellow
  - “Sonnet 18: Shall I compare thee to a summer’s day?” by William Shakespeare
  - “I Wandered Lonely as a Cloud” by William Wordsworth
  - “To Autumn” by John Keats
- rickastley.txt: Lyrics of “Never Gonna Give You Up” by Rick Astley
- unhr.txt: The Universal Declaration of Human Rights

These are available at `https://github.com/shubhamchandak94/nanopore_dna_storage_data/tree/master/encoded_file`. The files were tarred and compressed with bzip2 and then encrypted to randomize the data and avoid homopolymers. The final file size was 11,280 bytes. Note that while we use a 2 bits per base mapping in this work to convert the binary encoded data to the base sequences, previous works [7, 12, 13] have explored other mappings with desirable properties such as lack of homopolymers (e.g., GGGG) which are quite important when working with the basecalled reads due to systematic and biased errors in basecalling. In fact, an experiment in [12] showed that the consensus-based decoding failed when such a mapping was not used. Since we are not directly working with the basecalled sequence, we use the simple encoding that reduces the writing cost and makes the basecaller-decoder integration more straightforward, and rely on compression and encryption to avoid excessive homopolymers.

| m | r | RS redundancy | #oligos data | #oligos RS | #oligos total | Payload bits per oligo | Convolutional code input length | Oligo length | Writing cost (bases/bit) | #reads for decoding | Reading cost (bases/bit) |
|---|---|---|---|---|---|---|---|---|---|---|---|
| 8 | 1/2 | 30% | 1128 | 338 | 1466 | 80 | 100 | 108 | 0.91 | 5500 | 6.58 |
| 8 | 3/4 | 30% | 627 | 188 | 815 | 144 | 164 | 111 | 0.91 | 8000 | 10.20 |
| 8 | 5/6 | 30% | 564 | 169 | 733 | 160 | 180 | 114 | 1.75 | 13500 | 16.91 |
| 11 | 1/2 | 30% | 1128 | 338 | 1466 | 80 | 100 | 115 | 1.04 | 4500 | 5.54 |
| 11 | 3/4 | 30% | 627 | 188 | 815 | 144 | 164 | 117 | 0.92 | 6000 | 7.78 |
| 11 | 5/6 | 30% | 564 | 169 | 733 | 160 | 180 | 119 | 1.80 | 5500 | 7.01 |
| 14 | 1/2 | 30% | 1128 | 338 | 1466 | 80 | 100 | 113 | 1.06 | 3500 | 4.42 |
| 14 | 3/4 | 30% | 627 | 188 | 815 | 144 | 164 | 115 | 0.93 | 5000 | 6.59 |
| 14 | 5/6 | 30% | 564 | 169 | 733 | 160 | 181 | 117 | 1.85 | 6500 | 8.43 |
| 11 | 3/4 | 20% | 627 | 125 | 752 | 144 | 164 | 117 | 1.07 | 5500 | 7.13 |
| 11 | 3/4 | 40% | 627 | 250 | 877 | 144 | 164 | 117 | 0.95 | 5000 | 6.48 |

Table 1: Parameters for the experiments. The input to the convolutional code contains the index (12 bits) and the CRC (8 bits) in addition to the payload bits. For $m = 14, r = 5/6$ a padding of 1 bit was applied due to the boundary condition for the puncturing pattern. The oligo length and the writing/reading costs exclude the primer with length 25 bases on either end. The number of reads for decoding shown here correspond to list size of 8 for $m = 8, 11$ and list size of 4 for $m = 14$.

The input file was encoded into oligos for several experiments, as shown in Table 1. We varied the convolutional code memory $m$ and the rate $r$, and also tested a few values of the RS outer code redundancy for $m = 11, r = 3/4$. Additional encoding parameters such as the primers and the code used for encoding are available in `encode_experiments.py`. Note that the writing cost is calculated as (file size in bytes×8)/(#oligos total× Oligo length) while the reading cost is calculated as (file size in bytes×8)/(#reads for decoding×Oligo length).

#### 5.2 Main results

From the sequenced data, the raw signal files corresponding to each experiment were first separated as described in Section 6. A random sample of raw signal files per experiment was then decoded with the

| $m$ | $r$ | RS redundancy | Writing cost (bases/bit) | Reading cost (bases/bit) | | | | | | |
|---|---|---|---|---|---|---|---|---|---|---|
| | | | | $L=1$ | $L=2$ | $L=4$ | $L=8$ | $L=16$ | $L=32$ | $L=64$ |
| 8 | 1/2 | 30% | 1.75 | 7.18 | 7.18 | 6.58 | 6.58 | 5.98 | 6.58 | 6.58 |
| 8 | 3/4 | 30% | 1.04 | 12.74 | 11.47 | 11.47 | 10.20 | 10.83 | 12.11 | 14.02 |
| 8 | 5/6 | 30% | 0.92 | 20.66 | 18.78 | 18.16 | 16.90 | 19.41 | 21.29 | 25.04 |
| 11 | 1/2 | 30% | 1.80 | 6.15 | 6.15 | 5.54 | 5.54 | - | - | - |
| 11 | 3/4 | 30% | 1.06 | 9.08 | 7.78 | 7.78 | 7.78 | - | - | - |
| 11 | 5/6 | 30% | 0.93 | 8.92 | 8.28 | 7.01 | 7.01 | - | - | - |
| 14 | 1/2 | 30% | 1.85 | 5.05 | 5.05 | 4.42 | - | - | - | - |
| 14 | 3/4 | 30% | 1.07 | 7.25 | 7.25 | 6.59 | - | - | - | - |
| 14 | 5/6 | 30% | 0.95 | 9.08 | 9.08 | 8.43 | - | - | - | - |
| 11 | 3/4 | 20% | 0.98 | 8.43 | 7.78 | 7.78 | 7.13 | - | - | - |
| 11 | 3/4 | 40% | 1.14 | 7.78 | 7.13 | 6.48 | 6.48 | - | - | - |

Table 2: Reading costs for different list sizes.

convolutional code decoding (using `generate_decoded_lists.py`). The number of decoded reads was 20,000 for experiments with $m = 8$ and 10,000 for the rest (due to computational constraints). The list size was set to 64 for $m = 8$, 8 for $m = 11$ and 4 for $m = 14$. Note that the results for any list size lower than these can also be obtained by truncating the lists. This was followed by RS decoding of these lists for different list sizes using `decode_RS_from_decoded_lists.py`. For each list size, the minimum number of reads (in steps of 500) needed for successful decoding in 10 out of 10 subsampling trials was obtained and was used to compute the reading costs shown in Table 2. Based on this, the default list size of 8 was chosen for $m = 8, 11$ and 4 for $m = 14$.

From Table 2, we see that the reading cost typically reduces with the list size until a list size of 8, but then it starts increasing again. This is because the probability of having a random message in the list satisfying the CRC increases with the list size and hence the number of incorrectly decoded messages increases. This phenomenon is further explored in Table 3 which shows (for each experiment and list size), the percentage of reads that were (i) decoded correctly, (ii) had no CRC match in the list and, (iii) had a CRC match in the list that was incorrect. This was obtained using `compute_error_rate_from_decoded_lists.py`. Firstly, note that the percentage of correctly decoded reads increases with the list size, decreases with the code rate and increases with the convolutional code memory. This percentage is around 60%-80% for the $r = 1/2$ codes, around 30-40% for the $r = 3/4$ codes and around 20-30% for the $r = 5/6$ codes. As the list size increases to 64 for the $m = 8, r = 3/4$ code, the percentage of incorrect CRC matches grows as high as 9.33%, which means that around 18% of the RS redundancy is used up for the error correction leaving only $\sim$10% for recovering missing oligos, which leads to higher reading costs in Table 2.

#### 5.3 Impact of Reed Solomon outer code redundancy

Based on the analysis in [9], we expect that increasing the outer code redundancy should increase the writing cost and decrease the reading cost. Looking at the results in Table 2 for $m = 11, r = 3/4$ with RS redundancy 20%, 30% and 40%, we see that the reading costs for 40% reduncancy are the lowest which is as expected from the analysis. Interestingly, the 20% redundancy code has lower reading cost than the 30% redundancy code, but that might be due to randomness in the experiments and due to the higher number of correctly decoded reads for the 20% redundancy code (Table 3).

#### 5.4 Results for previous works

We compared the results for our approach with two previous works [7, 12]. The works in [7] and [12] use the same encoding based on Reed Solomon outer coding and constrained coding to avoid homopolymers, but the decoder in [12] uses an improved consensus algorithm leading to significantly lower reading cost. The writing cost for these works was computed as the reciprocal of the metric “bits per base excluding primers” reported as 1.10 in [7]. Since both these works report coverage (36x and 22x, respectively) rather than reading cost,

| | $L$ | 1 | 2 | 4 | 8 | 16 | 32 | 64 |
|---|---|---|---|---|---|---|---|---|
| $m = 8, r = 1/2$ | % correct | 59.65% | 63.35% | 66.33% | 68.93% | 70.99% | 72.57% | 73.98% |
| | % no CRC match | 40.26% | 36.49% | 33.35% | 30.47% | 27.79% | 25.40% | 22.40% |
| RS redundancy 30% | % incorrect CRC match | 0.06% | 0.13% | 0.29% | 0.58% | 1.20% | 2.01% | 3.60% |
| $m = 8, r = 3/4$ | % correct | 19.85% | 22.52% | 24.86% | 26.90% | 28.72% | 30.23% | 31.65% |
| | % no CRC match | 79.75% | 76.95% | 74.34% | 71.70% | 68.77% | 65.01% | 60.05% |
| RS redundancy 30% | % incorrect CRC match | 0.19% | 0.33% | 0.59% | 1.19% | 2.30% | 4.55% | 8.09% |
| $m = 8, r = 5/6$ | % correct | 15.40% | 17.90% | 19.98% | 21.67% | 23.53% | 25.12% | 26.54% |
| | % no CRC match | 84.27% | 81.61% | 79.25% | 76.87% | 73.75% | 69.85% | 63.96% |
| RS redundancy 30% | % incorrect CRC match | 0.17% | 0.33% | 0.61% | 1.30% | 2.55% | 4.87% | 9.33% |
| $m = 11, r = 1/2$ | % correct | 57.66% | 59.91% | 61.51% | 62.80% | - | - | - |
| | % no CRC match | 42.23% | 39.94% | 38.21% | 36.60% | - | - | - |
| RS redundancy 30% | % incorrect CRC match | 0.05% | 0.09% | 0.22% | 0.54% | - | - | - |
| $m = 11, r = 3/4$ | % correct | 31.47% | 34.87% | 37.42% | 39.62% | - | - | - |
| | % no CRC match | 68.25% | 64.66% | 61.95% | 59.30% | - | - | - |
| RS redundancy 30% | % incorrect CRC match | 0.15% | 0.34% | 0.50% | 0.95% | - | - | - |
| $m = 11, r = 5/6$ | % correct | 19.72% | 22.39% | 24.27% | 25.91% | - | - | - |
| | % no CRC match | 79.91% | 77.05% | 74.73% | 72.47% | - | - | - |
| RS redundancy 30% | % incorrect CRC match | 0.21% | 0.40% | 0.84% | 1.46% | - | - | - |
| $m = 14, r = 1/2$ | % correct | 76.49% | 78.66% | 80.15% | - | - | - | - |
| | % no CRC match | 23.39% | 21.18% | 19.60% | - | - | - | - |
| RS redundancy 30% | % incorrect CRC match | 0.06% | 0.10% | 0.19% | - | - | - | - |
| $m = 14, r = 3/4$ | % correct | 39.23% | 42.02% | 44.17% | - | - | - | - |
| | % no CRC match | 60.42% | 57.55% | 55.16% | - | - | - | - |
| RS redundancy 30% | % incorrect CRC match | 0.15% | 0.34% | 0.50% | - | - | - | - |
| $m = 14, r = 5/6$ | % correct | 24.23% | 26.40% | 28.24% | - | - | - | - |
| | % no CRC match | 75.43% | 73.05% | 70.88% | - | - | - | - |
| RS redundancy 30% | % incorrect CRC match | 0.15% | 0.36% | 0.69% | - | - | - | - |
| $m = 11, r = 3/4$ | % correct | 35.94% | 39.48% | 42.03% | 44.30% | - | - | - |
| | % no CRC match | 63.90% | 60.26% | 57.38% | 54.63% | - | - | - |
| RS redundancy 20% | % incorrect CRC match | 0.1% | 0.21% | 0.54% | 1.02% | - | - | - |
| $m = 11, r = 3/4$ | % correct | 29.82% | 32.77% | 35.14% | 37.14% | - | - | - |
| | % no CRC match | 70.03% | 66.98% | 64.40% | 61.77% | - | - | - |
| RS redundancy 40% | % incorrect CRC match | 0.09% | 0.19% | 0.40% | 1.03% | - | - | - |

Table 3: Percentage of reads correctly decoded, decoded with no CRC match found in list and incorrectly decoded with CRC match found. These add up to slightly below 100% since primer removal failed and convolutional code decoding was not performed for a very small percentage of reads (typically less than 0.2%).

we use the formula “reading cost = coverage × writing cost” for comparison with our results.

### 6 Experimental procedure

Oligonucleotides for all experiments were ordered as a single stranded DNA pool from GenScript. Pools were individually amplified using PCR primers specific to the experiment or subpool of interest using 1X Kapa HiFi PCR ReadyMix (Roche Biosystems), 500 nM PCR primers, and 1 microliter of the oligonucleotide pool under the following conditions: 98°C for 45 seconds; 12 cycles of 98°C for 15 seconds, 60°C for 30 seconds, and 72°C for 30 seconds; and a final extension step of 72°C for one minute. Each PCR reaction was purified with Ampure XP beads (Beckman Coulter) using standard protocols.

The thirteen pools were then quantified by fluorescence using the Qubit instrument (Thermo Fisher Scientific). They were then pooled and 300 femtomoles were used for downstream nanopore sequencing library preparation. Briefly, we performed End Repair and A-tailing of DNA products using the KAPA HyperPrep kit (Kapa Biosystems/Roche). The DNA was purified using Ampure XP beads (Beckman Coulter) under standard conditions. We then used the standard SQK-LSK109 library preparation from Oxford Nanopore Technologies to create DNA compatible for nanopore sequencing. Adapters containing motor protein was ligated to the DNA using 1X LNB buffer, AMX adapter, and KAPA HyperPrep ligase enzyme. The library was again purified using Ampure XP beads, but washed with SFB buffer and eluted in EB buffer (both Oxford Nanopore Technologies). The libraries were then loaded and sequenced on a MinION 9.4.1 flowcell for 48 hours using standard protocols.

After synthesis and sequencing, the reads were basecalled using the default Guppy basecaller (by ONT) and aligned to the original sequences sent for synthesis. These reads were then separated into the respective experiments and the corresponding raw signal data was used to perform the decoding. Note that we used alignment to separate the various experiments for simplicity, but they can be separated either physically using PCR or computationally using the distinct primers. To enable comparison across experiments, only the reads that align to the original sequences were considered for decoding. In practice, the unaligned/low-quality reads could be filtered out based on the primer sequence or using the CRC. The script `util/align_compute_stats.sh` was used for these steps.

### 7 Basecalling analysis

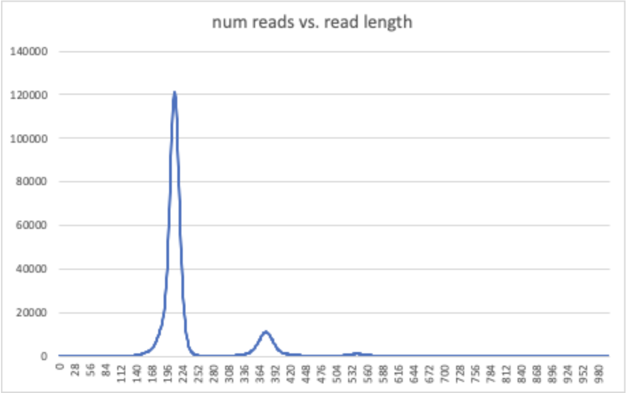

Figure 3: Read length distribution of the basecalled reads. There is a small peak at twice the expected length indicating chimeric reads due to ligation.

Even though the proposed approach works directly on the soft information provided by the basecaller and not on the basecalled sequences, we performed some analysis on the basecalled reads for completeness. The basecalled reads had close to 10.5% error rate with around 3-4% each of insertions, deletions and substitutions. Most of the errors are due to nanopore sequencing, since Customarray synthesis error rate is below 1% based on previous work in [9]. Out of a total of 3.4 million sequenced reads, around 30%

were unaligned and around 8% were chimeric (i.e., multiple oligos ligated together - see Figure 3 for the read length distribution). The code in `util/read_length_distribution.cpp` was used for finding the read length distribution.

While the experiments were underway, a new high accuracy version of Guppy basecaller was released. We performed basecalling with this and the alignment was better, leading to only 15% unaligned reads with average basecalled error rate of 8.4%. The error rate is still quite high and does not affect our main conclusions regarding the efficacy of basecaller-decoder integration. Note that the Flappie basecaller used for the decoding of the raw signal already uses the high accuracy model, so the main results concerning the reading cost for this approach are independent of this update.
